# Examining the Ensembles of Amyloid-*β* Monomer Variants and their Propensities to Form Fibers Using an Energy Landscape Visualization Method

**DOI:** 10.1101/2021.10.28.466349

**Authors:** Murilo N. Sanches, Kaitlin Knapp, Antonio B. Oliveira, Peter G. Wolynes, José N. Onuchic, Vitor B. P. Leite

**Affiliations:** Department of Physics, São Paulo State University (UNESP), Institute of Biosciences, Humanities and Exact Sciences, São José do Rio Preto, SP 15054-000, Brazil; Center for Theoretical Biological Physics, Rice University, Houston, TX 77005, USA; Departments of Physics and Astronomy, Chemistry, and Biosciences, Rice University, Houston, TX 77005, USA

## Abstract

The amyloid-*β* (A*β*) monomer, an intrinsically disordered peptide, is produced by the cleavage of the amyloid precursor protein, leading to A*β*40 and A*β*42 as major products. These two isoforms generate pathological aggregates, whose accumulation correlates with Alzheimer’s disease (AD). Experiments have shown that even though the natural abundance of A*β*42 is smaller than that for A*β*40, the A*β*42 is more aggregation-prone compared to A*β*40. Moreover, several single-point mutations are associated with early-onset forms of AD. This work analyzes coarse-grained AWSEM simulations of normal A*β*40 and A*β*42 monomers, along with six single-point mutations associated with early on set disease. We analyzed the simulations using the Energy Landscape Visualization Method (ELViM), a reaction coordinate-free approach suited to explore the frustrated energy landscapes of intrinsically disordered proteins. ELViM is shown to distinguish the monomer ensembles of variants that rapidly form fibers from those that do not form fibers as readily. It also delineates the amino-acid contacts characterizing each ensemble. The results shed light on the potential of ELViM to probe intrinsically disordered proteins.

## Introduction

Most of molecular biology has been guided by the structure-function paradigm, whereby the three-dimensional structure of a protein determines specific biological functions . The process by which a newly created peptide sequence can achieve an organized structure can be quite complex, but is now understood by considering the energy landscape theory,^1–5^ More recently, a broad class of proteins whose amino acid sequences result in only minimally structured proteins has been recognized. Commonly referred to as intrinsically disordered proteins (IDPs),^6,7^ these peptides appear to explore rough, relatively flat energy landscapes that are marked by a multitude of local basins having similar free energ ies.^8–10^ These proteins lack structure when fully solvated, but may undergo large conformational changes to more ordered states under certain conditions.^11,12^ Yet, until such triggering conditions are met, they explore a large conformational space.^13–15^ Some IDPs while disordered tend to exist in relatively compacted conformations,^16^ which are the result of competition between entrop y and energy. Computer simulations have provid ed insight into the conformational transitions they can undergo. A major problem in such studies is how to analyz e and classify the large and highly variable data sets that are produced.^17^

A significant number of neurological diseases have been associated with IDPs.^18,19^ Often, IDP monomers can interact to form aggregate structures, rang ing over a large scale of sizes, which may either still be disordered or may be highly ordered. Determining the mechanisms driving the aggregation process from individual unstructured monomers to structured fibrils is critical to understanding the role of aggregates in disease. ^20^ Over the last 20 years, a huge number of research efforts have been directed toward understanding the aggregation processes associated with Alzheimer’s Disease (AD), a form of dementia suffered by an estimated 50 million people worldwide.^21^ One of the proteins closely associated with AD is a peptide fragment known as amyloid-beta (A*β*). The most abundant forms of A*β* are either 40 or 42 residues in length.^22^ The apparent neural damage in AD has long been thought to be directly caused by the presence of large fibers and plaques,^23^ but small, soluble oligomers of A*β* ^23,24^ have also been considered the source of the damage. Understanding A*β* in its monomeric form is a useful place to start in any case. Two regions in the monomers have been identified as playing key roles in the dynamics of aggregation, the central hydrophobic core (CHC), consisting of residues 17-20, and the B-turn region formed by residues 23-28. The B-turn region is capable of forming a salt-bridge, which has been observed to accelerate the aggregation process.^25^ Despite differing by only two amino acids in length, A*β*-40 and A*β*-42 markedly differ in their aggregation tendencies. While A*β*-40 is naturally present at a ratio of 10:1 over A*β*-42 in cerebrospinal fluid, A*β*-40 is only present in the much smaller ratio of 1:3 in amyloid plaques as compared to A*β*-42,^26^ consistent with the experimental observation that the nucleation and elongation rates of A*β*-42 are much faster than those of A*β*-40.^27^ Studies have previously shown that the behavior of these isoforms diverge early in the aggregation process during the formation of small oligomers.^28^ The aggregation rate for A*β*-42 correlates with the *β*-strand content found in its monomeric form.^29^ It should also be noted that several single-point mutantations of amyloid-*β* have been identified^30^ that are associated with early onset and inherent forms of the disease.^31–33^

To investigate the correlation of the monomer structural ensembles with the aggregation propensities of A*β*, we have used the Energy Landscape Visualisation Method (ELViM) ^34^ to analyze the conformational ensembles of A*β* 40 and 42 monomers, as well as the landscapes of six A*β*-40 mutations. The ELViM technique has already exhibited a capacity to discriminate and clarify the energy landscapes of IDPs, as shown in a previous work with the prostate associated gene 4 (PAGE4).^35^ Through the local structural similarity metric proposed by Wolynes et. al.,^36^ it is possible to survey and triangulate a high-dimensional conformational phase space, and then using ELVIM, project the ensembles to two optimal dimensions while preserving local proximities. This allows an intuitive visual analysis of the energy landscape.

## Methods

### Simulations Details

Choosing a force field to simulate a free monomer of A*β*, in both its 40 and 42 residue variants, is crucial. The ideal force field for modeling disordered peptides should have a strong physical foundation, incorporate information from ideal geometries and known electrostatic properties, and be shown to make accurate structure predictions. For this reason we used the Associative-memory, Water-mediated, Structure and Energy Model (AWSEM)^37^ force field. This coarse-grained force field, was obtained by using energy landscape based machine learning, run in LAMMPS,^29^ and can explore a large energy landscape. The A*β*-42 simulations were initiated starting from a solution structure of A*β*-42 (1Z0Q) inferred from NMR.^30^ In the case of A*β*-40 the decision was made to use the same starting structure, though truncated to 40 residues. Comparable solution studies of A*β*-40 (2LFM)^31^ also show significant local helical secondary structure and therefore were started from the same conditions. Each simulation was initialized by energy minimization with a 2fs timestep for the first 100,000 timesteps, followed by a simulation run of 10^6^ timesteps. During this run, structures were sampled every 1000 timesteps. Simulations were carried out at 250K, 300K, 350K, and 400K, with two independent simulations per specified temperature. This resulted in a total of 0.4 µs of simulation time, and 100,000 sampled structures. Further details regarding these simulations can be found in the Supplemental Information section. For the mutant A*β*-40 species, simulations were also started from the same annealed structure to avoid any secondary structure bias, since the structures of the monomeric mutants have yet to be resolved. The simulation procedure was consistent with that previously described, with the data from the full range of temperatures used to determine the energy landscape, though only simulations carried out a 300K were used for structural sampling and comparison.

### Energy Landscape Visualisation Method

For globular folded proteins, conformational changes can usually be represented in terms of an effective reaction coordinate associated with a unique native state. This approach is not as useful for IDPs because their landscapes are not strongly funneled. To visualize the details of the energy landscape of A*β*, we instead employ the Energy Landscape Visualization Method (ELViM). This method starts with a metric based on the internal distances between pairs of amino acids.^34^ One measure of similarity between two conformations *k* and *l* that is quite useful is given by:

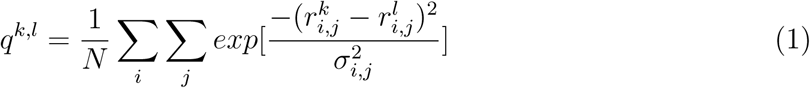

where *N* is the total number of pairs of residues,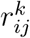 is the distance between the residue *i* and *j* in the conformation *k*, 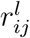 is the distance in the conformation *l*and *σ*_*i,j*_ is a weighing constant equals to *σ*_0_|*i - j*|^*ε*^, with *σ*_0_ = 1 and *ε* = 0.15. The normalized distance between the conformations can then be defined by *δ*^*k,l*^ = 1 *- q*^*k,l*^.

The methodology is divided into three steps: (I) The first step to create a visualization of the landscape is to calculate the distances between every pair of structures in the trajectory, building a *nXn* dissimilarity matrix. (II) The second step consist of a data clustering based in the distance between the conformations in the matrix. Structures close to each other by less than a given cut-off distance are clustered into a single conformation. (III) The last step is the multidimensional reduction of the original data to a 2D or 3D projection using these clusters as a guide. A multidimensional projection technique initially places the objects of the matrix at an random initial positions in a 2 or 3-dimensional space and then applies a minimization step to preserve the distances in the dissimilarity matrix. The process stops when the difference in distances between consecutive iterations is below a given cut-off.

For the projections in this work we used the simulated monomers of A*β*-40 and A*β*-42 at the 250K, 300K and 350K temperature, forming the conformational phase space, and the mutants structures at 300K.

## Results and Discussion

Since A*β* is an intrinsically disordered protein, describing conformational changes would require more than one reaction coordinate. ELViM is a reaction-coordinate free method that allows one to visualize in an optimal way the high-dimensional conformational phase space of each type of A*β* in a small number of dimensions. By analysing all the sampled conformations at once, we can make a comparison of the conformational ensembles of different studied monomers. We create a large projection with the A*β*-40, A*β*-42, and A*β*-mutants conformations, all represented and analyzed in a single phase-space. Figure 1 shows this projection with the different species separated in different colors in (A) and the entire projection labelled in colors indicating the radius of gyration in (B). Each conformation of the data-set is represented by a point mapped in the 2D projection. One can see that most of the A*β*-40 points are located on the right side of the projection, while the A*β*-42 points are more widely spaced, and the mutants follow a similar behavior. The structures around the projection in Figure 1.B exemplifies the conformations of each region, which shows that the regions most populated by A*β*-40 correspond to the *α*-helix clusters, which are also more compact. In contrast, the regions with a bigger radius of gyration correspond to the A*β*-42 and A*β*-mutants.

**Figure 1:**
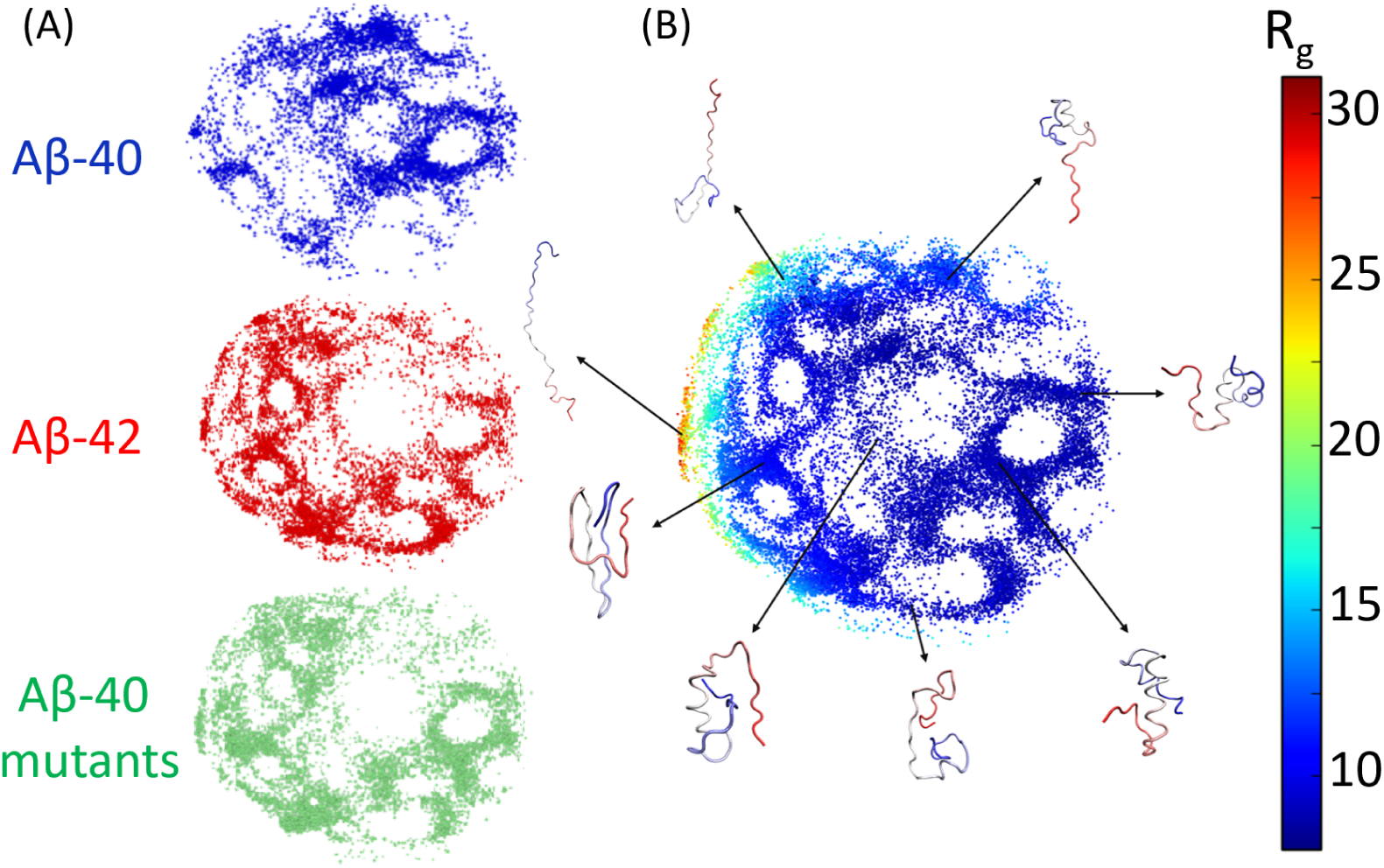
Conformational phase space of the simulated structures visualized through ELViM.(A)Distinct species present in the projection, with the A*β*-40 in blue, A*β*-42 in red, and the A*β*-40 mutants in green. (B) Each conformation as function of the radius of gyration, with the most significant conformations o each region displayed around it.

This result is supported by Figure 2, which shows the ELViM projection as a function of the *q* coordinate measured with respect to the A*β*-40 monomer (2LMN) in the fiber as reference (*q*_*fiber*_). We see that the A*β*-40 landscape populates a region with low *q*_*fiber*_, while A*β*-42 and the mutants landscapes populate a region with somewhat high *q*_*fiber*_. This suggest a reason for the much higher propensity for fiber formation by A*β*42 and the mutants.

**Figure 2:**
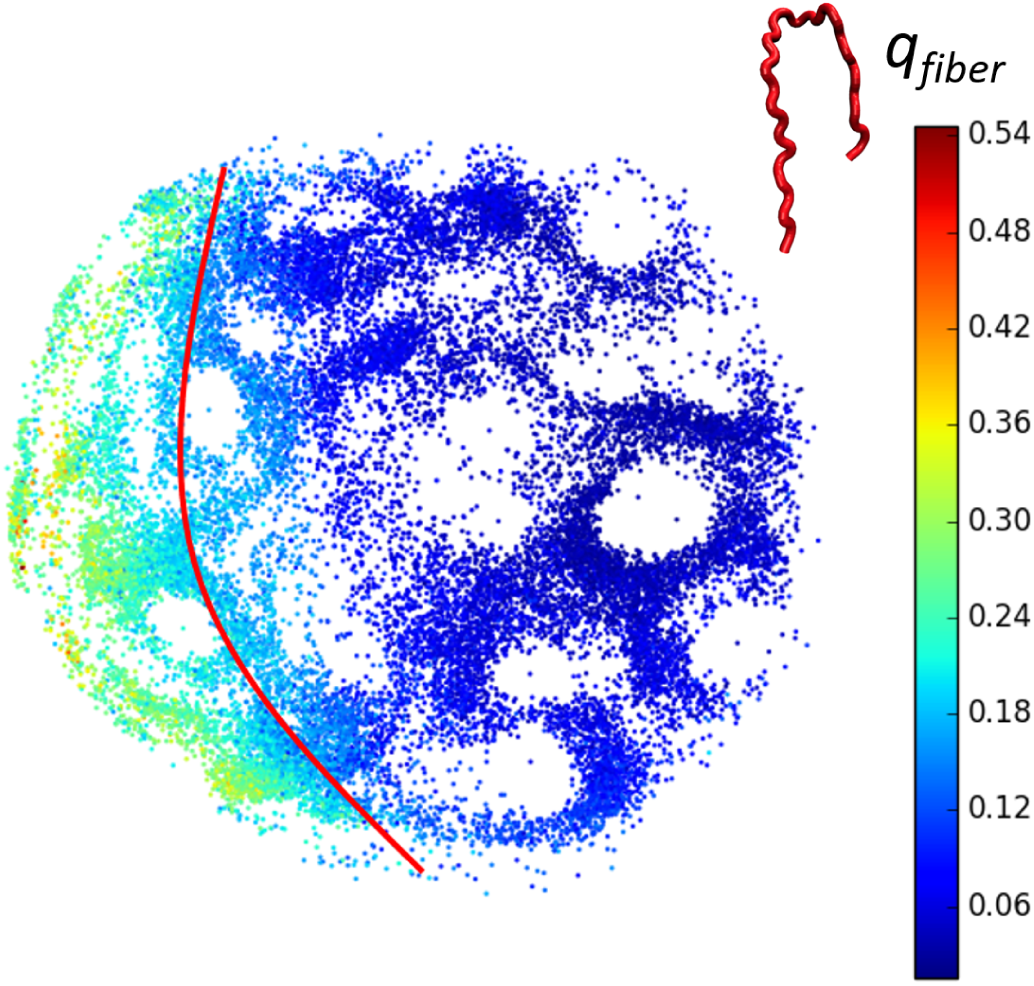
2-dimensional ELViM projection as a function of *q* with the fiber monomer as reference (red structure, PDB 2LMN). The red line act as a divider between a high and low *q*_*fiber*_ region in the projection.

To examine this finding more closely, using the Weighted Histogram Analysis Method (WHAM), we calculated the free energy as a function of the similarity to the fiber monomer structure. The results shown in Figure 3 suggest there is an energetic difference between extended and more fiber-like structures for the A*β*-40, which is less prominent for the A*β*-42 and the mutants. A*β*-40 presents an almost continuously increasing free energy as a function of increasing *q*_*fiber*_, while the other systems seem to have a more two-state-like behavior. Table 1 shows the values for the free energy profiles, with the Δ*F* ^*‡*^ indicating the valor of the energetic barrier, while the Δ*F* indicates the height of the first free energy minimum state. Despite having high free energy cost for fiber like ordering, the D23N mutation behave like the A*β*-42, as will be discussed below.

**Figure 3:**
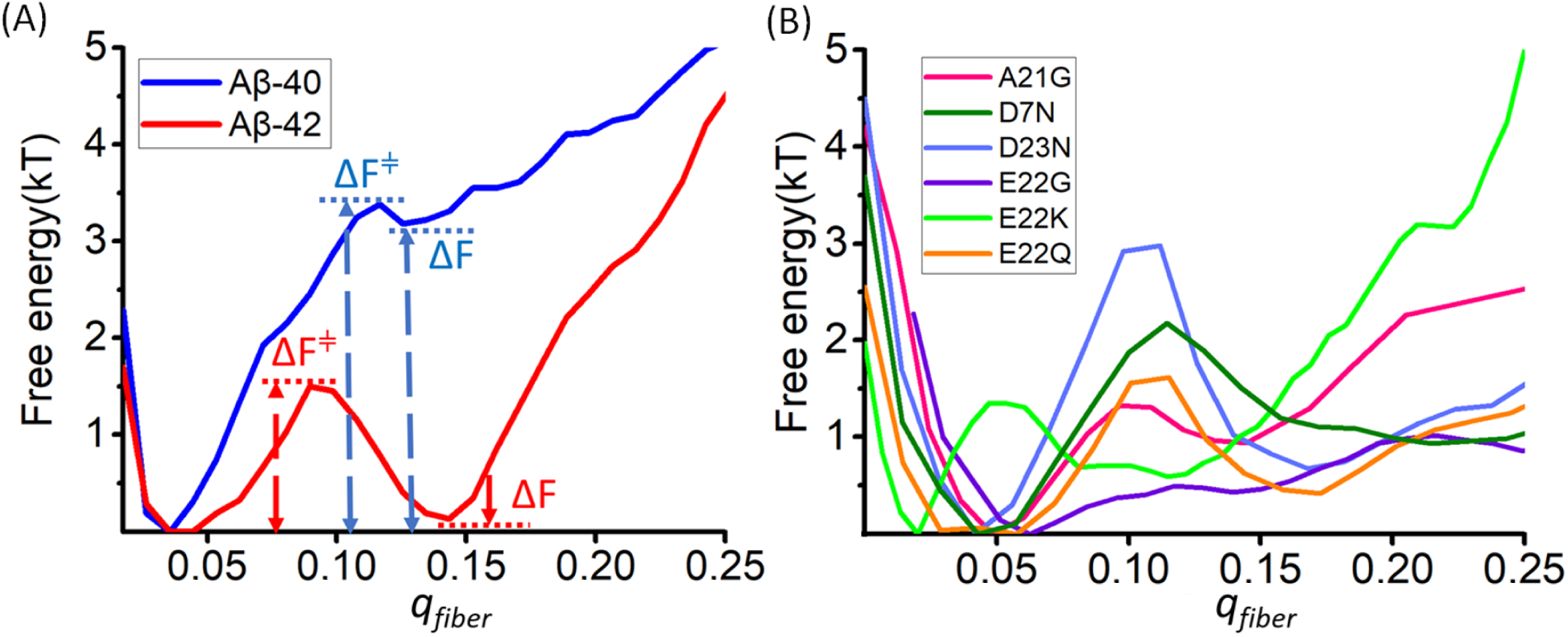
(A) Free energy profile as a function of *q*_*fiber*_ for the A*β*-40 and A*β*-42. (B) Free energy profile as a function of *q*_*fiber*_ for the A*β*-40 mutants.

**Table 1:** Free energies

| Species | $\Delta F(\text{kT})$ | $\Delta F^\ddagger(\text{kT})$ |
| --- | --- | --- |
| $A\beta$-40 | 3.2 | 3.7 |
| $A\beta$-42 | 0.2 | 1.6 |
| A21G | 0.9 | 1.3 |
| D7N | 0.9 | 2.1 |
| D23N | 0.9 | 2.9 |
| E22G | 0.8 | 0.9 |
| E22K | 1.1 | 2.5 |
| E22Q | 0.4 | 1.5 |

We next analyze the distribution of the structures in the ELViM projection, by reweighing the conformations using the density of microstates Ω(*E*) obtained from the WHAM.^38^ Figure 4 shows the result in a grid-like projection with each bin’s color indicating the free energy of the ensemble of structures contained within. The region of the *α*-helix in the Figure 1 corresponds to the lowest free energy structural cluster for the A*β*-40, as is pointed out with the red *G* in the upper part of the Figure 4.A, while the more fiber-like region identified by Fig. 4.B is highlighted with the black ellipse. The red and yellow lines indicate proposed routes for reconfiguration (the lowest free energy path) when A*β*-40 transverses to assume the fiber-like conformations, with Figure 5 illustrating the most probable conformational changes along each path. The lower part of 4.A exhibits the grid projection for the A*β*-42, in which the free energy profile indicates that the lowest free energy structures are nearer the fiber region than for the A*β*-40 case. The most likely pathway of reconfiguration (yellow line) is shorter and more direct, implying that the most stable conformations already have some of the contacts contacts necessary to form a fiber. Figure 6 displays the most representative structures of this path. Video animations showing the conformational changes along the possible routes of Figures 5 and 6 are shown in the SI. The locations of the lowest free energy for both isoforms confirm that the free energy gap between these states and the aggregation-prone states is smaller for A*β*-42 than for A*β*-40, agreeing with Figure 3. This suggest that in some way the aggregation propensity for each peptide may already be encoded in the monomer free energy spectrum.^39^ The same can be seen for theA*β*-40 mutants in Figure 7, with all low free energy states located in the same region as A*β*-42 and, as the yellow paths show, following the behavior similar to A*β*-42.

**Figure 4:**
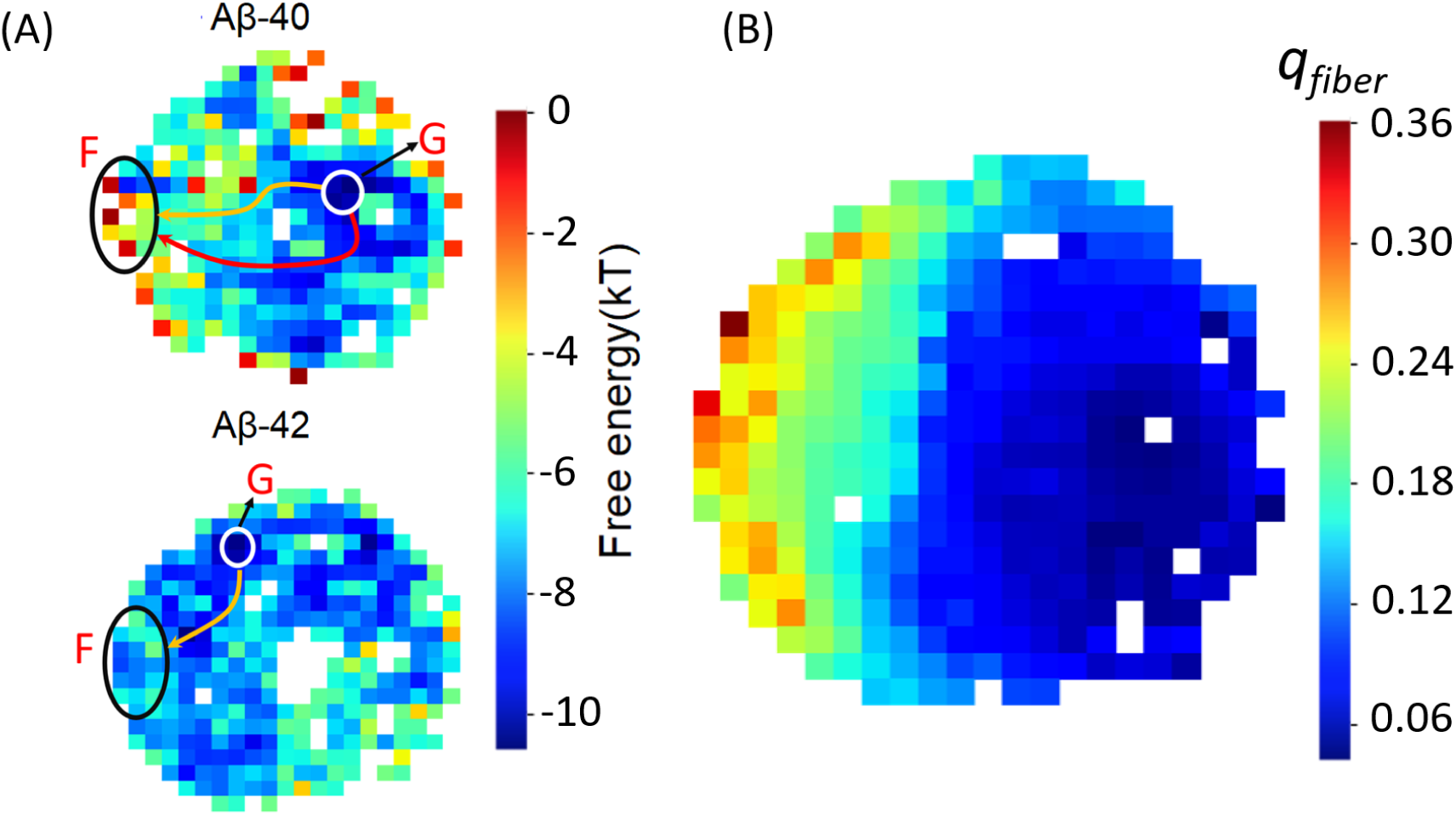
(A) Free energy distribution of the A*β*-40 (upper) and A*β*-42 (lower) structures in the ELViM projections, with the lowest region indicated by the G and the region of high *q*_*fiber*_ indicated by the ellipse with the F. (B) Mean *q*_*fiber*_ distribution of the Figure 2.

**Figure 5:**
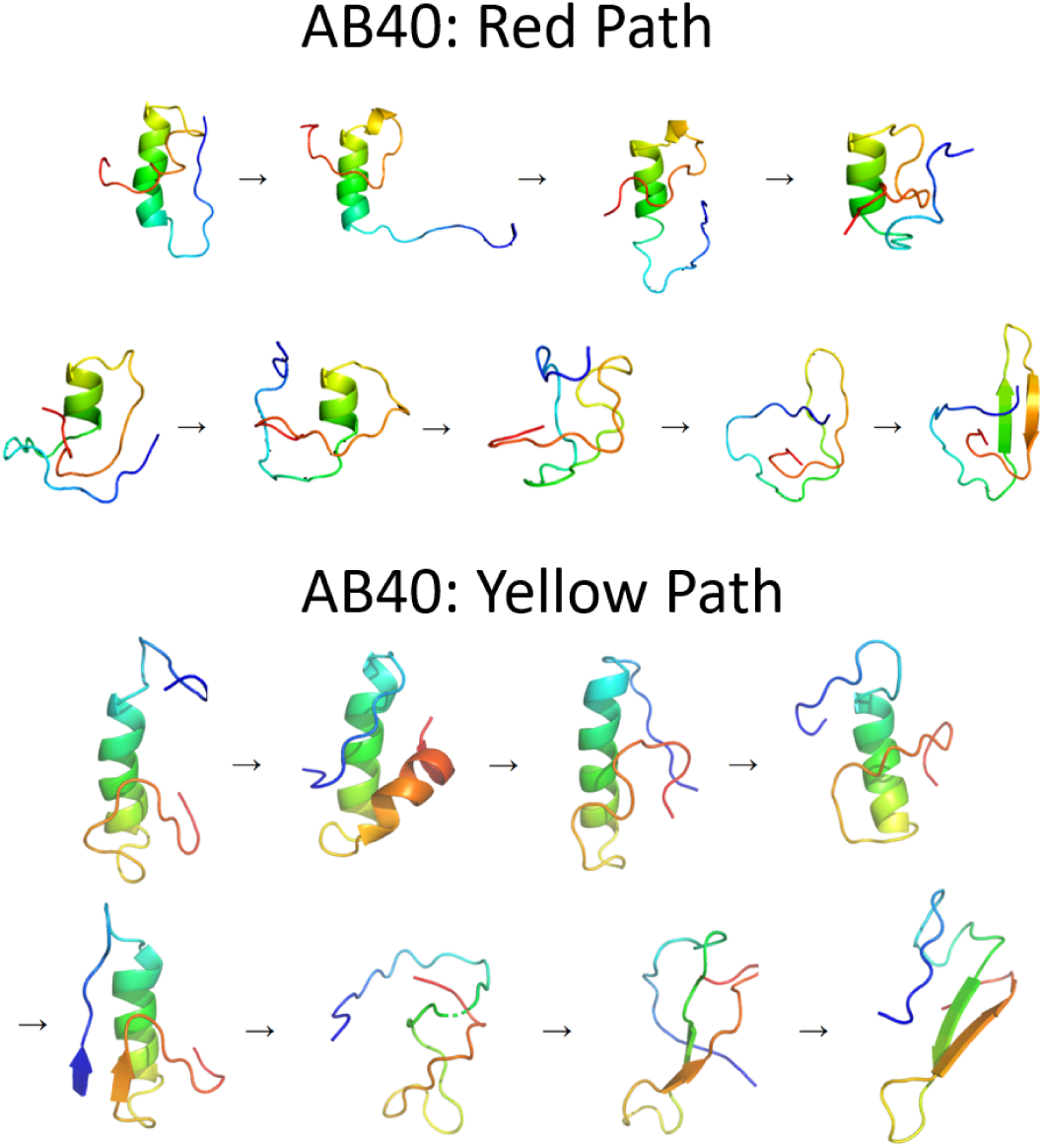
Conformational changes of the structures contained in the possible paths of A*β*40 to morph into fiber-like structures.

**Figure 6:**
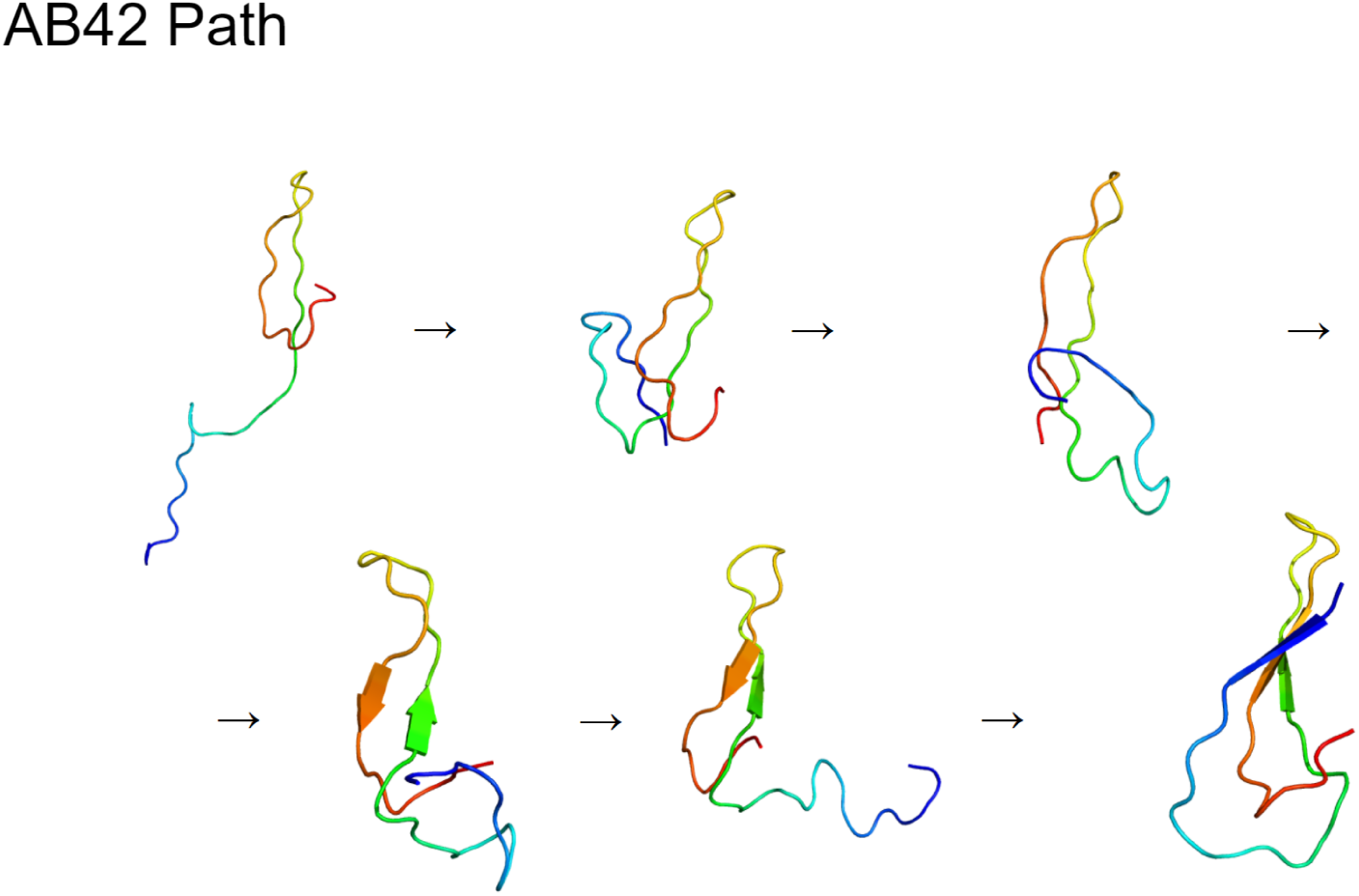
Conformational changes of the structures contained in the possible path of A*β*42 to morph into fiber-like structures.

**Figure 7:**
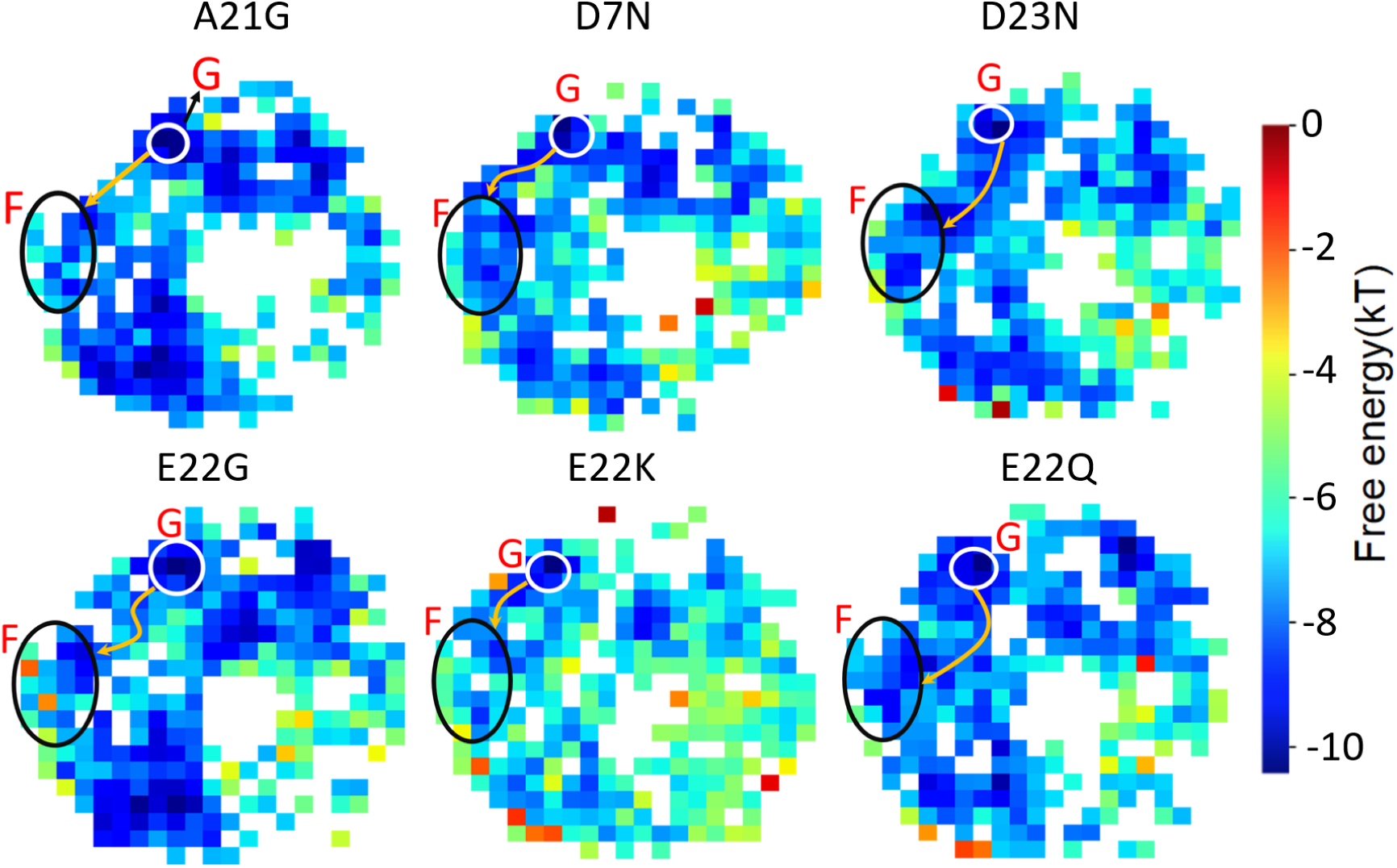
Free energy distribution for each A*β*-40 mutation with the pathway of lowest free energy highlighted in yellow.

Based on this observation, we calculated the differences in the contact maps averaged over the structures located in the lowest free energy bin and those averaged over structures in the fiber region. For this analysis, we removed the last two residues from A*β*-42, so we could compare the maps between the species. Figure 8 shows these differences for the A*β*-40, 42 and the mutation E22Q. Apparently, the A*β*-40 molecule must first undo the *α*-helix to begin the formation of the first *β*-sheet contacts, while A*β*-42 instead undergoes a chain-sliding mechanism, visible in the parallel contacts between residues 15 to 30, to achieve the fiber-like structure. The E22Q mutation does not have the entire *β*-sheet formed, but also does not exhibit the *α*-helix contacts, and thus does not need to overcome a high energetic barrier to form the fiber, as is seen in Figure 3. The contact maps, along with the ELViM projections, confirm that the A*β*-42 has a higher propensity to form *β*-strands than A*β*-40 and its mutants, in harmony with earlier reports.^40^ The differences in the contact maps for the others mutations are available in the supplementary information in Figure S1, along with the contact maps for each species (Figure S2).

**Figure 8:**
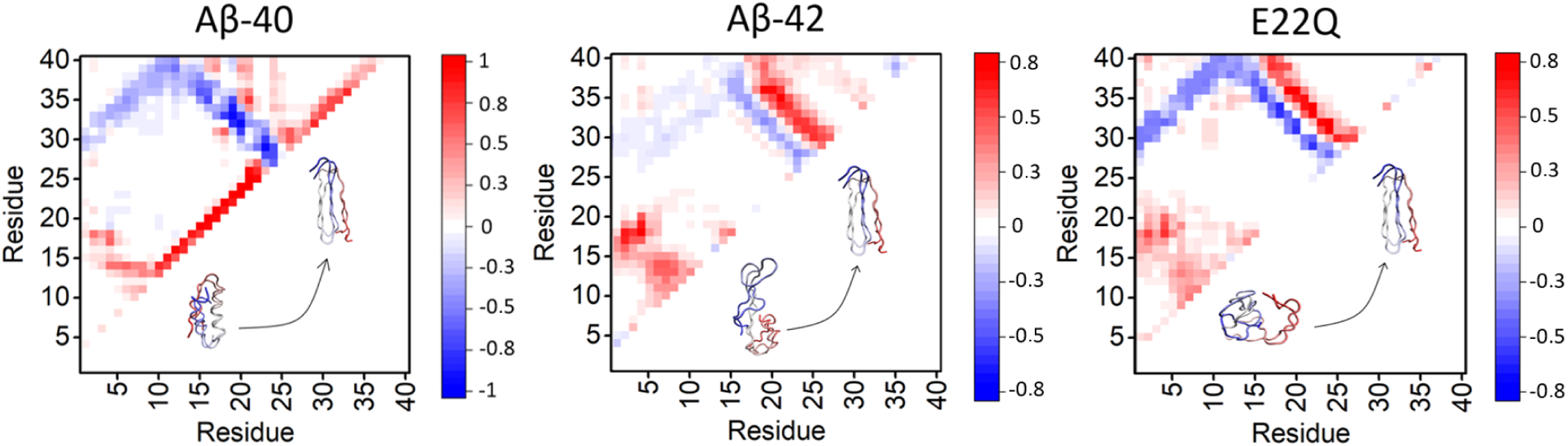
Contact maps. Difference between the mean contact maps for the regions of lowest free energy (red/positive values) and those from the regions of high *q*_*fiber*_ (blue/negative values) for the A*β*-40, A*β*-42 and E22Q structures.

## Conclusion

The disordered nature and correspondingly large structural fluctuations of IDPs lead to a complex high dimensional energy landscape. The lack of a funnel to single dominant structure poses a significant challenge for analysis. Unlike for globular protein folding, it is difficult to determine adequate reaction coordinates. Nevertheless, ELViM allows one to visualiz e the high dimensional phase space in an easy-to-understand 2D projection, without the need for a native conformation or a specific reaction coordinate. It is thus a powerful tool for the study of IDPs. In the present work, ELViM permitted the exploration of the conformational changes in A*β* in its monomeric forms. This study suggests the aggregation rate is correlated with the *β*-strand content of the monomers, since the A*β*-42 and the mutants have higher aggregation rates and exhibit a lower free energy cost to form *β*-sheet than the more soluble A*β*-40. Moreover, the low free energy regions in the projection of the ensembles show that most of the contacts necessary to start the aggregation have already been formed in the the A*β*-42 monomers and the monomers of the less soluble mutants, while the monomers of A*β*-40 need to undo some helical structure first.

## Acknowledgement

MNS was supported by Conselho Nacional de Desenvolvimento Científico e Tecnológico (CNPq, Grant 130147/2020-6). This research was supported by the Center for Theoretical Biological Physics sponsored by the NSF (Grants PHY-2019745). JNO is also supported by the NSF (Grant CHE-1614101) and by the Welch Foundation (Grant C-1792). JNO is a Cancer Prevention and Research Institute of Texas (CPRIT) Scholar in Cancer Research. ABOJ acknowledges the Robert A. Welch Postdoctoral Fellow program. PGW is also supported by the D.R. Bullard-Welch Chair at Rice University (Grant C-0016). VBPL was supported by CNPq (Grant 310017/2020-3) and FAPESP (Grants 2019/22540-3 and 2018/18668-1).

