## Supplemental Information for "Examining the Ensembles of Amyloid-*β* Monomer Variants and their Propensities to Form Fibers Using an Energy Landscape Visualization Method"

**Video S1.** Video A40red.mp4 shows the A$\beta$40 conformational changes to morph from the lowest free energy region to fiber-like structures through the **red** path shown in Figure 5.a.

**Video S2.** Video Ab40yellow.mp4 shows the A$\beta$40 conformational changes to morph from the lowest free energy region to fiber-like structures through the **yellow** path shown in Figure 5.b.

**Video S3.** Video Ab42.mov shows the A$\beta$40 conformational changes to morph from the lowest free energy region to fiber-like structures through the path shown in Figure 6.

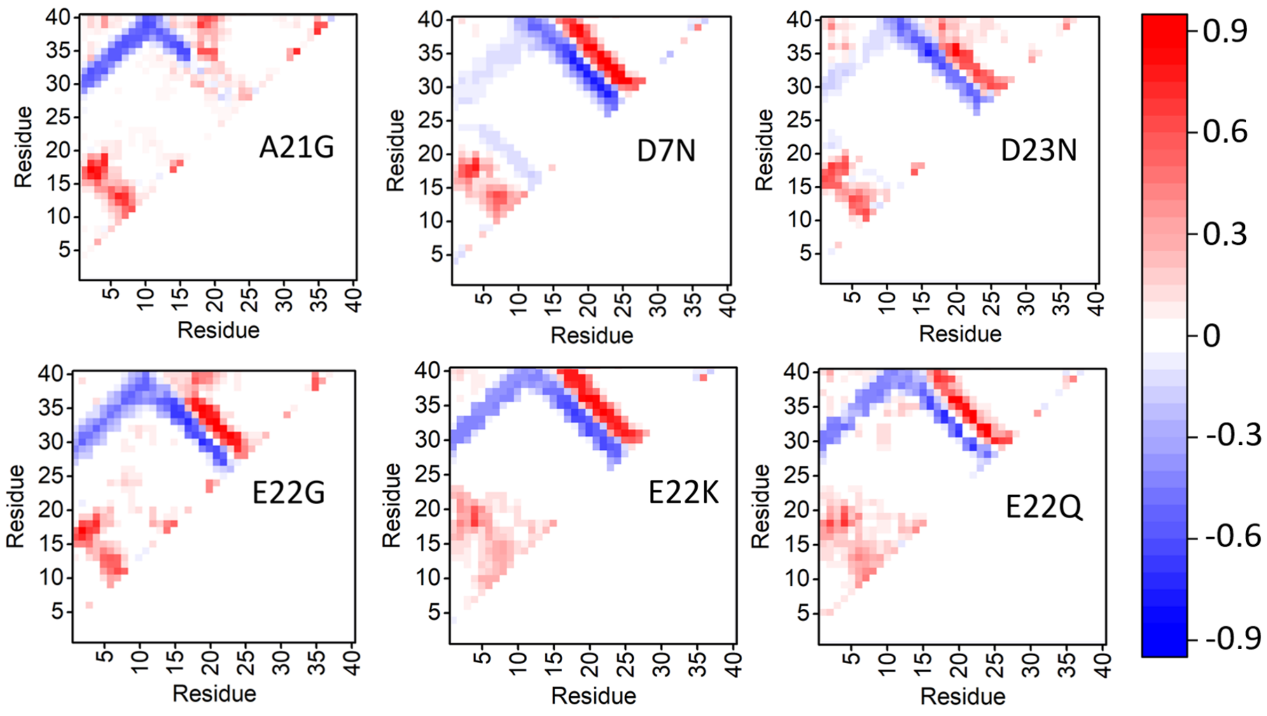


Figure S1: Difference between the mean contact maps for the regions of lowest free energy with the regions of high $q_{fiber}$ for each A$\beta$-40 mutant.

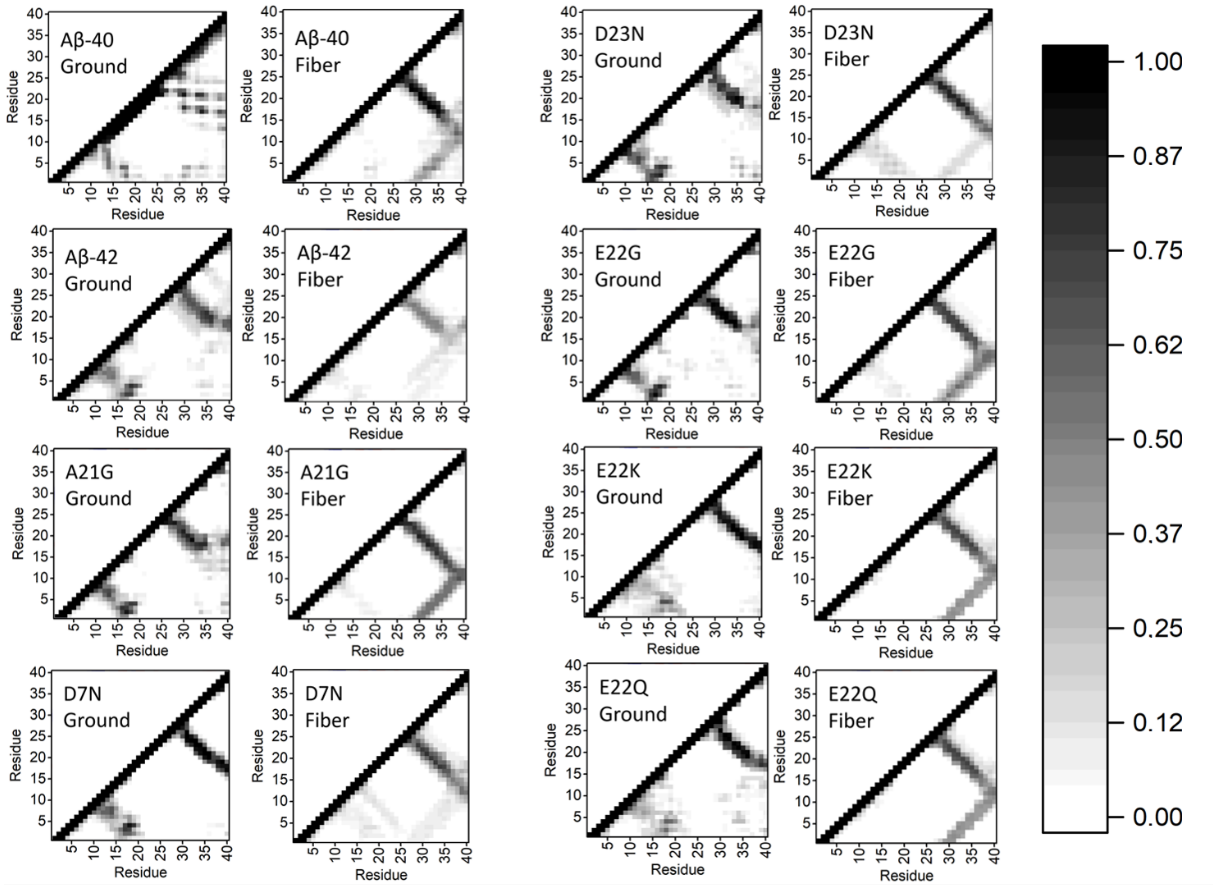


Figure S2: Mean contact maps for the regions of the lowest free energy (ground state) and the fiber-like structures (high $q_{fiber}$) for each A$\beta$-40 mutant.
